# Extending the limits of 3D printed polymers on paper towards bioanalytical sensing

**DOI:** 10.64898/2026.03.27.714910

**Authors:** P. Ngaju, R. Pandey, K. Kim

## Abstract

Polymeric 3D printing of microfluidic devices for biosensing is an appealing fabrication alternative for rapid manufacturing of biosensing devices with complex geometry in a streamlined, repeatable and cost-effective manner without the need for expensive instrumentation such as those employed in photochemical etching and soft lithography. Hybrid 3D printed paper-based microfluidics is an emerging area which harnesses the unique properties of both, merging the construction of microfluidic structures and the inherent capillary-driven flow within paper substrates. In this work, we have fabricated hydrophobic barriers by 3D printing a single layer of machinable wax, thermoplastic polyurethane, polylactic acid and polypropylene directly on chromatography paper to create open microchannels and determine the most suitable material. Characterization of each open microchannel using the four materials revealed polypropylene as the most reliable material with high hydrophobic barrier integrity and resolution. Polypropylene achieved functional microchannels with a resolution of 621 ± 33µm, hydrophobic barrier integrity of (93.75 ± 9.16%), wicking speed of 0.38mm/s and optimal hydrophilicity of channels (51.4 ± 8.36 °) with minimal embedding during thermal curing. To demonstrate proof of principle, a fluorescence assay demonstrating the formation of a dimeric g-quadruplex structure from a g-rich sequence which significantly enhances fluorescence of thioflavin T was implemented.

## Introduction

Microfluidic paper-based analytical devices (µPADs), a subfield, harnesses the unique properties and advantages of cellulose based paper as an inexpensive pump-free analytical platform which include porosity, inherent capillarity, biocompatibility with biological samples, biodegradability, ease of modification^1,2^. The Whitesides’ research group was the first to report µPADs in 2007, designing a 3D-µPAD with multiple stacked layers. Despite this achievement, photolithography was used as the fabrication method to pattern the paper which is a costly process requiring access to a clean room and a series of complicated steps^3^. µPADs have since been used in numerous applications such as bioassays, environmental monitoring and blood tests employing various detection methods such as colorimetry, fluorescence, electrochemical, chemiluminescence among others^4–7^. Hydrophobic barriers have been traditionally created using wax and a commercial printer to print 2D patterns created using CorelDraw or open-source vector graphics software such as Inkscape^8^. Despite this simple, low cost approach, there are several limitations with using wax, some of these include the discontinuation of wax printers such as the Xerox ColorQube, the lack of flexibility with the thickness of chromatography paper that can be used, constraints with channel width resolution due to difficulties controlling the embedding of wax during the thermal curing process and barrier integrity issues when certain surfactants and organic solvents are used hence limiting the scope of bioanalytical applications that can be used with wax barriers^9^. A handful of new studies have published the possibility of 3D printing hydrophobic barriers, notably, Suvanasuthi et al. showed the feasibility of using wax filament to print hydrophobic barriers for point of care detection of dengue virus serotypes^10^ and Ng et al. used a combination of polycaprolactone to create hydrophobic barriers on filter paper and polylactic acid for 3D structures^11^. Furthermore, He et al., 3D printed polylactic acid (PLA) channels, sealed them with polydimethylsiloxane (PDMS) and filled the channels with cellulose powder for capillary-driven flow^12^. 3D printing directly on filter paper was reported by Zargaryan et al. to create hydrophobic barriers, finger-operated reservoirs and reversible mechanical valves to enable fluid control without the use of pumps, however, no results were reported on the highest functional channel resolution (smallest channel width that can allow efficient wicking of fluids) or an application for biosensing^13^. Despite these advancements in the fabrication of 3D printed paper-based devices, biosensing demonstrations using this approach are limited. There is still room to create simple, robust and repeatable fabrication methodologies with high channel resolution for the purposes of bioanalytical fluorescence based assays. Here we report a simple two-step fabrication process of directly 3D printing a single layer of material on chromatography paper using machinable wax, polypropylene (PP), thermoplastic polyurethane (TPU) and PLA, optimizing 3D printing parameters, comparing and contrasting strengths of each with respect to degree of material embedding after thermal curing using scanning electron microscopy, determining hydrophobic barrier integrity and wicking efficiency. PP emerged as the most reliable material and was further characterized using contact angle measurements within the paper hydrophilic channels and polypropylene barriers. Furthermore, the effect of nozzle size on channel resolution was investigated and finally the approach was validated with a lateral flow PP 3D µPAD device validated with a dimeric g-quadruplex (G4-dimer) formation assay. Dimeric g-quadruplexes are higher order nucleic acid structures in which two individual g-quadruplex scaffolds stack or associate to form a joint four-stranded architecture stabilized by G-tetrads and associated cations such as sodium ions ^14,15^. G-quadruplex scaffolds strongly bind to small molecule fluorophores such as thioflavin T (ThT) which then undergo a large fluorescence enhancement with the interaction with the dimeric structure^16^. In biosensing, dimeric G-quadruplexes are important as fluorescent signal-amplification elements by embedding in DNA probes or amplification products so that the formation of the dimeric g-quadruplex structure leads to a pronounced turn on fluorescence without the need for covalent labeling of the recognition sequence^17^. The transformation of a single stranded dimeric g-quadruplex sequence into an anti parallel dimeric g-quadruplex structure is achieved in the presence of high concentrations of sodium ions and binding with a cationic benzothiazole dye, ThT, significantly enhanced fluorescence intensity.

## Experimental section

### Materials and chemicals

Tris-EDTA ethylenediaminetetraacetic acid (HO_2_CCH_2_)_2_NCH_2_CH_2_N(CH_2_CO_2_H)_2_) (TE) buffer solution at pH 8.0, tris hydrochloride(tris-HCl), sodium chloride (NaCl), thioflavin T, ultrapure DNase/RNase-free distilled (MilliQ) water were all purchased from Millipore Sigma (Vermont, US). The oligonucleotides used in this study namely, dimeric g-quadruplex (G4-Dimer) (5′-GGG TAG GGC GGG TTG GGG GGT AGG GCG GGT TGG G-3′) and a control oligonucleotide (5’ TTT CTT GGA TGG TGA TGC ATG GCC GTT TTT AGT TCG TGA ATA TCG TAT TTG CCG CTA A 3’) were purchased from Integrated DNA Technologies (Iowa, US). Print2Cast/machinable wax filament was purchased from MachinableWax.com Inc (Michigan, US), Polypropylene (PP), Thermoplastic polyurethane (TPU) and polylactic acid (PLA) filaments were obtained from Shop3D (Mississauga, ON). Whatman Grade 1 cellulose chromatography paper with a thickness of 0.18mm and a linear flow rate of 130mm/30min was purchased from Cytiva Life Sciences (Massachusetts, US).

### Apparatus and software

3D patterns were designed using SolidWorks Academic Version (Autodesk). A fused deposition modeling (FDM) 3D printer (Qidi Tech X-Plus II) (China) was used to 3D print hydrophobic barriers directly on chromatography paper. A Phenom Pro X benchtop scanning electron microscope with energy-dispersive X-ray spectroscopy was used to characterize the structure and morphology of microchannels for devices created using wax, PP, TPU and PLA. Contact angle measurements were conducted using a Dataphysics OCA15EC optical contact angle & contour analysis instrument. To capture fluorescence images a Nikon Eclipse Ti2 microscope with a numerical aperture of 0.20, 4x PlanApo objective, episcopic resolution of 1.53µm, brightness of 1.0 and pixel size (µm/pixel) of 1.625 with an excitation wavelength range of 446-486 nm and emission wavelength range of 500-550 nm. Image J Fiji licensed under the GNU public licence version 2 was used post-image capture to quantify the degree of fluorescence in each image.

### Fabrication and characterization of 3D printed µPADs

Prior to 3D printing hydrophobic barriers, the heating bed of the 3D printer was levelled to accommodate for the thickness of the chromatography paper. Hydrophobic barriers were created by 3D printing structures one layer high, directly on a 25 x25 mm chromatography paper affixed to the heating bed with masking tape. 3D printing parameters were optimized based on each material. After 3D printing, the devices were thermally cured in a convection oven, cooled for 5 minutes and laminated with a lamination film at the bottom before using in experiments.

The 3D-printed µPAD channels were characterized using scanning electron microscopy to determine, surface morphology and channel porosity. Hydrophobic barriers fabricated using machinable wax, PP, TPU and PP were fabricated to determine the degree of embedding and the deviation between channel widths before and after thermal curing for 1000 µm and 700 µm devices (n = 10 discrete devices) and compared. Furthermore, hydrophobic barrier integrity was assessed for (n = 10 discrete devices) for each material by calculating the percentage of devices that reliably contained TE buffer coloured with food dye. Finally, wicking speed efficiency was studied and the best performing material was selected based on all comparative experiments. The effect of nozzle size on channel resolution was studied for the best performing material by 3D printing using 4 different nozzle sizes 0.2, 0.25, 0.3 and 0.35 mm. ThT was deposited within the channels and fluorescence images captured before using Image J Fiji to measure the channel widths. Three measurements were taken for each device (n=4 discrete devices) and thereafter analysed using one way ANOVA. Image J Fiji was used to measure the channel widths after thermal curing, thereby investigating the degree of embedding and the deviation between actual and measured channel widths. Contact angle experiments using deionized water were conducted on the best performing material to determine the degree of hydrophilicity of the channels in comparison to the barrier and determination of parameters for the Lucas Washburn equation that models the relationship between imbibition distance and time.

### Validation of dimeric g-quadruplex formation fluorescence assay in solution and on 3D printed µPAD

A solution assay was first conducted to verify the formation of a dimeric g-quadruplex structure from a g-rich sequence before implementation on the 3D-µPAD. A 100 µM stock solution of the g-rich sequence was prepared in 5 mM Tris-HCl (pH 7.4), heated at 90°C for 10 min and then slowly cooled to room temperature, thereafter diluted to the desired concentrations (10 µM, 8 µM, 6 µM, 4 µM and 2 µM) in 150 mM NaCl buffer. A stock solution of ThT (1mM was prepared in deionized water (∼18.2 MΩ cm), diluted to 20µM and stored at 4 °C in the dark. 50 µl of the G4-Dimer was pipetted into aliquots for each of the five concentrations, thereafter 50 µl of 20 µM ThT and incubated in the dark for 90 min at 20 °C. A control oligonucleotide was prepared in an identical manner as the G4-Dimer, the negative control was 20 µM ThT. A BMG Clariostar microplate reader was used to measure the fluorescence intensity from each sample in triplicate. The excitation wavelength was set to 425 nm and the emission wavelength 485nm.

The paper-based G4-Dimer formation assay was implemented as signal ‘on/off’ where fluorescence intensity of the G4-Dimer was compared to that of the control oligonucleotide. The G4-Dimer sequence and the control oligonucleotide were diluted in equimolar concentrations to the desired concentrations (20 nM, 40 nM, 60 nM, 80 nM and 100 nM). 5 µM of ThT was deposited onto the 3D µPAD and allowed to dry in the dark for 24 hours before using in experiments. After running the assay with n=3 discrete devices for each concentration, fluorescence images of the sensing regions were captured with a Nikon Eclipse Ti2 microscope and fluorescence intensity quantified using Image J Fiji.

### Statistical Analysis

All measurements acquired from the experiments were plotted to show mean ± standard deviation. GraphPad Prism software version 10.5.0 was used for statistical analysis of all results. One-way and two-way ANOVA were used as the statistical tests considering a 95% confidence interval with Tukey’s and Sidak’s multiple comparisons tests.

## Results and Discussion

### Fabrication and characterization of 3D printed microfluidic devices

Hydrophobic barriers were created by directly 3D printing on chromatography paper one layer high with four materials: PLA, PP, TPU and wax as shown in ***Figure 1***. The heating bed was levelled to accommodate for the thickness of the chromatography paper (0.18mm). An infill density of 100% was selected to ensure even coating of the paper, furthermore, slower printing speeds including appropriate levelling of the heating bed ensured good adhesion of the polymer to the paper yielding reliable hydrophobic barriers. G-code settings for the printer were adjusted and optimized based on the material being printed and have been summarized in ***Table 1***.

**Table 1:** Optimized 3D printing parameters and thermal curing temperatures for wax, TPU, PP and PLA for the creation of hydrophobic barriers on chromatography paper.

| *Filament material* | *Nozzle temperature (°C)* | *Print bed temperature (°C)* | *Infill density (%)* | *Infill speed (mm/s)* | *Outer wall speed (mm/s)* | *Inner wall speed (mm/s)* | *Thermal curing temperature (°C)* | *Thermal curing duration (min)* |
| --- | --- | --- | --- | --- | --- | --- | --- | --- |
| Wax | 150 | 80 | 100 | 20 | 20 | 25 | 150 | 5 |
| TPU | 230 | 50 | 100 | 20 | 20 | 25 | 200 | 30 |
| PP | 220 | 80 | 100 | 30 | 20 | 25 | 200 | 27 |
| PLA | 210 | 60 | 100 | 30 | 20 | 25 | 205 | 30 |

**Figure 1.**
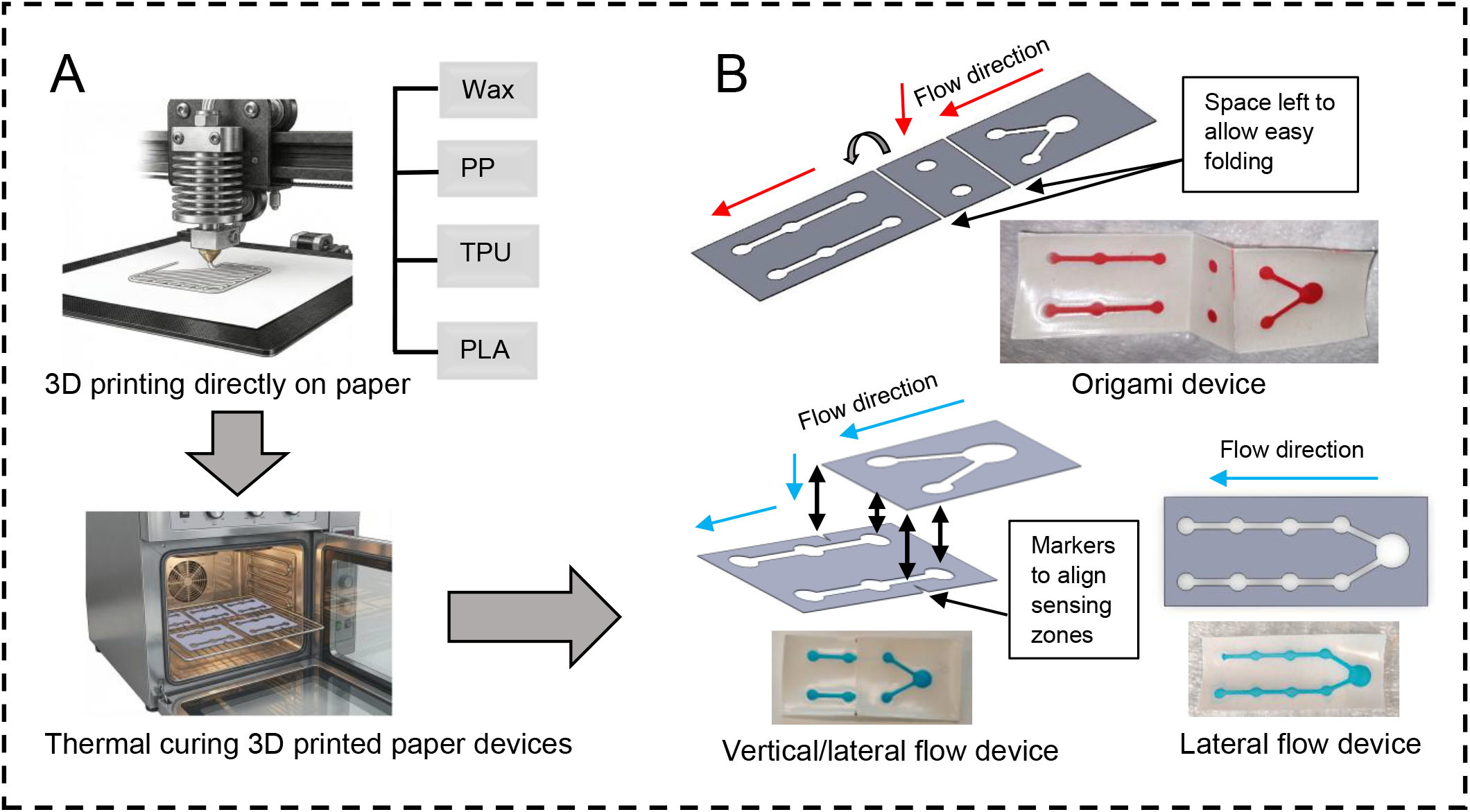
Schematic showing fabrication process of **A.** 3D printed paper devices **B**. versatility of the approach with three different capillary flow configurations with actual devices.

During thermal curing, high temperatures are used to allow the material to penetrate and embed within the paper, subsequently creating hydrophobic barriers. The fibres in paper are mostly anisotropic and tend to be more horizontal than vertical, therefore, lateral spreading of fluids is usually more rapid in comparison to vertical spreading when deposited material is heated^18^. The 3D printed devices were characterized using SEM to determine the surface morphology and degree of material embedding within the channels to assess wicking capacity as shown in ***Figure 2***. To better understand the changes in channel width before and after thermal curing, n=5 devices with channel widths of 700um and 1000um were designed and measured before and after heating (***Figure 2I&J***).

**Figure 2.**
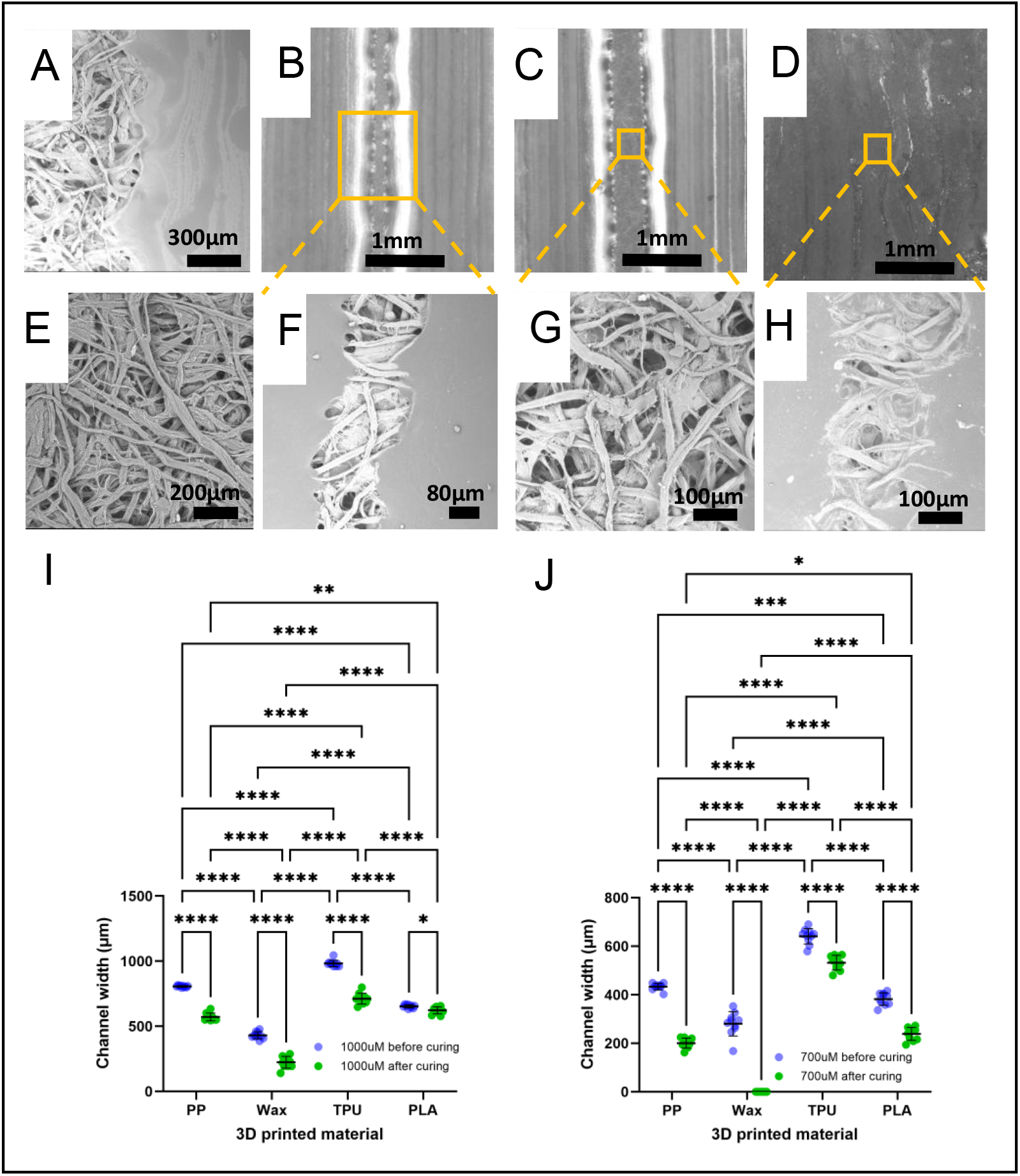
SEM images of paper device channels. (A) TPU (B,F) PP, (E) chromatography paper, PP, (C,G) PLA, (D,H) wax, I compares as fabricated to after thermal curing 1000 µm channel widths J compares as fabricated to after thermal curing 700 µm channel widths. *p<=0.05, **p<=0.01, ***p<=0.001, ****p<=0.0001.

TPU appeared to evenly coat the surface of the paper (***Figure 2A***), however penetration and embedding within the paper fibers was shallow leading to poor hydrophobic barrier integrity (33.33± 26.06%). PP exhibited robust hydrophobic barriers with clear visibility of the porous paper fibres, fully embedded hydrophobic barriers, minimal channel embedding (***Figure 2F***) and excellent hydrophobic barrier integrity (93.75 ± 9.16%) indicating deep penetration of material within the paper Fibres. Furthermore, PLA demonstrated high channel porosity similar to paper (***Figure 2E***), however embedding was poor leading to low barrier integrity (3.75±10.61%). Upon heating wax spreads laterally (in the x- and y-direction) as well as penetrating the paper fiber (z-direction) when heating leading to significantly more material embedding within the channels in comparison to PLA, PP and TPU (***Figure 2D***). As a result the porosity of the channels was low as shown in (***Figure 2H***). SEM results correlated with hydrophobic barrier experiments where channel widths below 1600µm were fully embedded and unusable but overall optimal barrier integrity was high (84 ± 30.50%) (***Table 2***). Each material was assessed to determine retention of the original channel width for 700 µm and 1000 µm devices (***Figure 2I&J***). For the 1000 µm, the as fabricated TPU channel width was closest to the original measured width. It is helpful to note that the material is extruded at high temperatures before it is deposited on the paper, therefore primary material embedding occurs when the material comes into contact with paper during 3D printing before post fabrication thermal curing, resulting in reduced channel widths (***Figure 2I***). PP is the second closest followed by PLA then wax. The channel width of PP compared to PLA is statistically significant (p<=0.001) with PLA showing minimal variation between as fabricated and thermally cured devices. A similar trend was observed for 700µm devices with TPU closest to the actual channel width followed by PP, PLA then wax (***Figure 2J***) with a slight difference in the statistical significance between PP and PLA (p < 0.01). Wax channels after thermal curing were fully embedded with no porosity observed to measure the channel width, results that corroborate SEM images in ***Figure 2H***.

**Table 2.** Hydrophobic integrity of materials shown as percentage of devices that contained 50µl of liquid without leaking(n=10 discrete devices were used to calculate %)

| Channel width ($\mu\text{m}$) | Material | | | |
| --- | --- | --- | --- | --- |
|  | TPU (%) | PLA (%) | Wax (%) | PP (%) |
| 1000 | 66.67 | 0.00 |  | 100.00 |
| 1200 | 10.00 | 30.00 |  | 100.00 |
| 1400 | 10.00 | 0.00 |  | 100.00 |
| 1600 | 0.00 | 0.00 | 30.00 | 80.00 |
| 1800 | 40.00 | 0.00 | 100.00 | 100.00 |
| 2000 | 70.00 | 0.00 | 100.00 | 90.00 |
| 2200 | 30.00 | 0.00 | 100.00 | 80.00 |
| 2400 | 40.00 | 0.00 | 90.00 | 100.00 |
| Mean | 33.33 | 3.75 | 84.00 | 93.75 |
| SD | 26.06 | 10.61 | 30.50 | 9.16 |

To assess and characterize the wicking efficiency, devices for each material were fabricated with two channel widths, 1000 µm and 1800 µm. Coloured food dye was added to Tris buffer and used to visualize and measure the time it took for the buffer to wick from one reaction pad to another within each channel for PLA, PP, PP and wax. Results summarized in ***Figure 3*** show an overall increase in wicking velocity as the channel size increases. The data shows that the integrity of the hydrophobic barriers has an impact on the wicking speed efficiency. From 1.0 mm to 1.6 mm PP the wicking speed is not significantly different, at 1.8mm there is a jump in the speed which is thereafter maintained for 2.0 - 2.4mm channel widths (***Figure 3I***). TPU showed a consistent increase in wicking speed with an increase in channel width with some leaking within the barriers. Below 1.6mm wax channels are fully embedded and cannot wick liquid (***Figure 3I***). A slight drop in wicking speed is observed from 1.6mm to 1.8mm, this observation is due to challenges with controlling wax’s isotropic flow upon melting and the paper’s anisotropic porosity leading to inconsistent hydrophobic barriers^18^. PLA had a similar trend compared to TPU, however a decline in speed between 1.4mm and 1.6mm was observed, which was attributed to challenges with hydrophobic barrier integrity.

**Figure 3:**
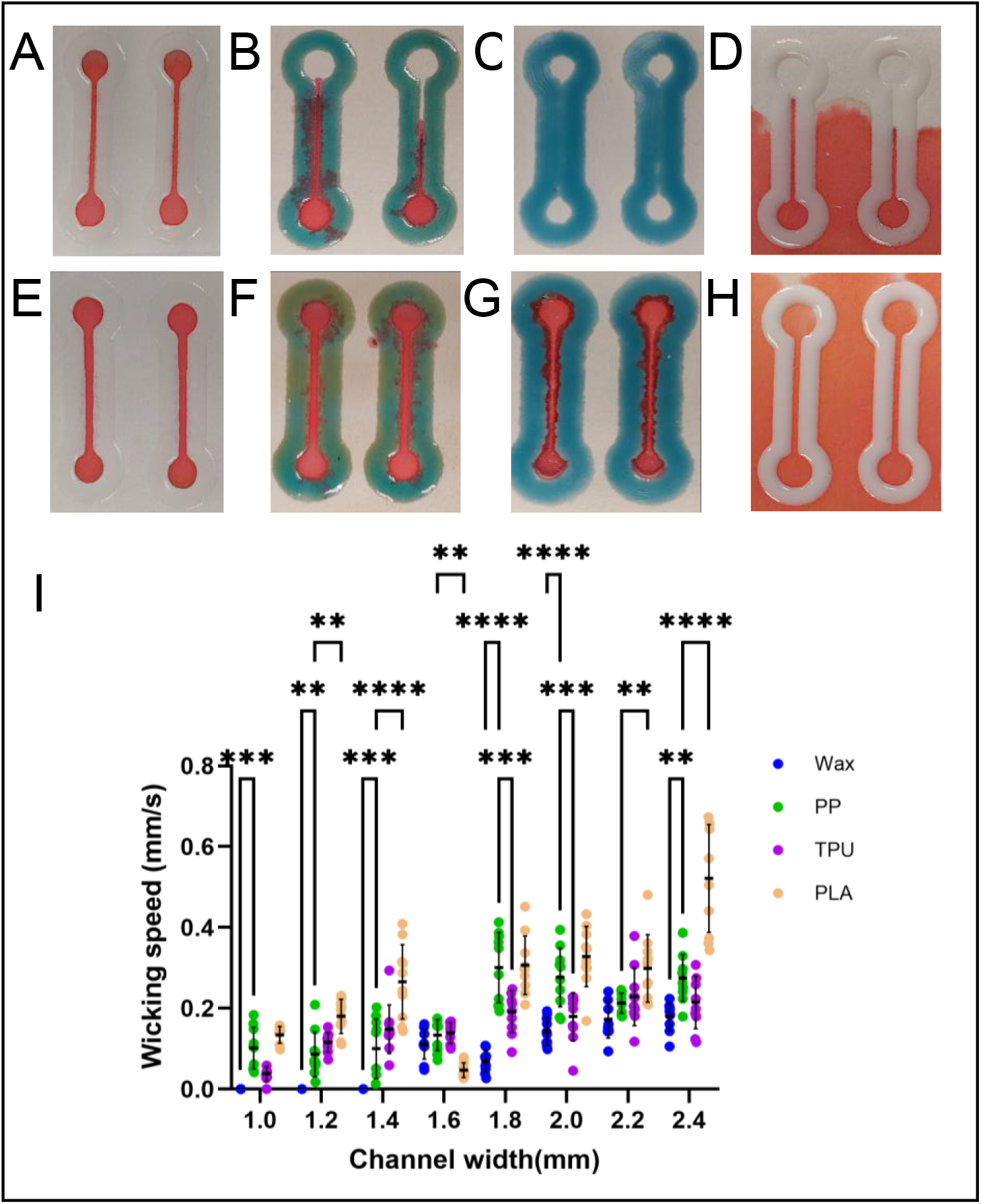
Wicking speed efficiency experiments for PLA, PP, TPU and wax were conducted by deposition 50µl of coloured food dye on each device. **A** 1000 µm PP devices B 1000 µm TPU devices **C** 1000 µm wax devices D 1000 µm PLA devices **E** 1800 µm PP devices F 1800 µm TPU devices **G** 1800 µm wax devices H 1800 µm PLA **devices I** wicking speed vs channel width. *p<=0.05, **p<=0.01, ***p<=0.001, ****p<=0.0001

From the results, PP emerged as the best performing polymer with optimal hydrophobic barrier integrity, wicking speed and minimal channel embedding post thermal curing. PP achieved microchannels with a high resolution of 621 ± 33µm (***Figure S2***). Furthermore, the versatility of PP was demonstrated in the design and fabrication of three device configurations, vertical lateral flow, origami and lateral flow devices as shown in ***Figure 1***. Using PP channels, the effect of 3D printing with different nozzle sizes was investigated. From the results (***Table S1***) there were no significant differences when channels 3D printed using 0.2mm was compared with 0.3mm including 0.2mm compared with 0.35mm nozzle sizes. However when comparing 0.2mm to 0.25mm there was a significant difference (p < 0.01) as well as between 0.25mm and 0.35mm (p < 0.01) as shown in ***Figure 4A&B***. The significant differences are attributed polypropylene embedding within the channels attributed to capillary-driven imbibition of the molten phase into the porous cellulose network that is modified by temperature-dependent viscosity, surface tension and the complex pore structure of the paper. Several factors could be responsible for significant variations in channel width for smaller nozzle sizes these include 1) variation with adhesion of the polymer during initial 3D printing of hydrophobic barriers, less material deposited could be responsible for poorer adhesion and 2) variations in the penetration depth during post printing thermal curing processes. These factors result in non-linear behaviour typical of non-Newtonian fluid flow due to variations in fluid viscosity caused by temperature gradients. Larger nozzle sizes (0.30 and 0.35mm), however, exhibit less variations, possibly due to sufficient deposition of material during 3D printing creating optimal adhesion with the paper leading to more robust channels reducing the standard deviation between the actual and measured channel widths. To compare hydrophilicity of the microchannels (***Figure S3***) and hydrophobicity of the barriers (***Figure S4***) contact angles of both were measured. Results revealed hydrophilic microchannels optimal for fluid wicking (51.4 ± 8.36 °) and barriers tending towards hydrophobicity at (82.6 ± 6.27 °). Furthermore, to characterize imbibition of liquid within the PP 3D µPAD channels, contact angles (***Figure S3***) were used to model capillary flow using the Lucas Washburn equation (***eq. (1)&(2)***) for open microchannels. The Lucas Washburn equation describes the progression of a fluid front to be directly proportional to the square root of the time that has passed since the initial wicking point contacted the fluid^19^. 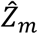 is the position of the meniscus as a function of time 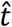 for flow in Newtonian liquids, 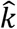 is the mobility parameter that can be described as the diffusion coefficient driving the growth of the liquid interface, *R*is the effective pore radius of the matrix which is usually approximated, the chromatography paper used had a pore radius of ∼10µm, *θ* is the contact angle (51.4 ±8.36 °for 1800µm PP 3D µPAD device), *µ* is the viscosity of the liquid and 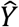 the surface tension of the liquid air interface^19^.

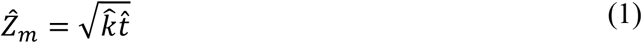

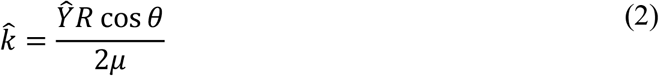

**Figure 4:**
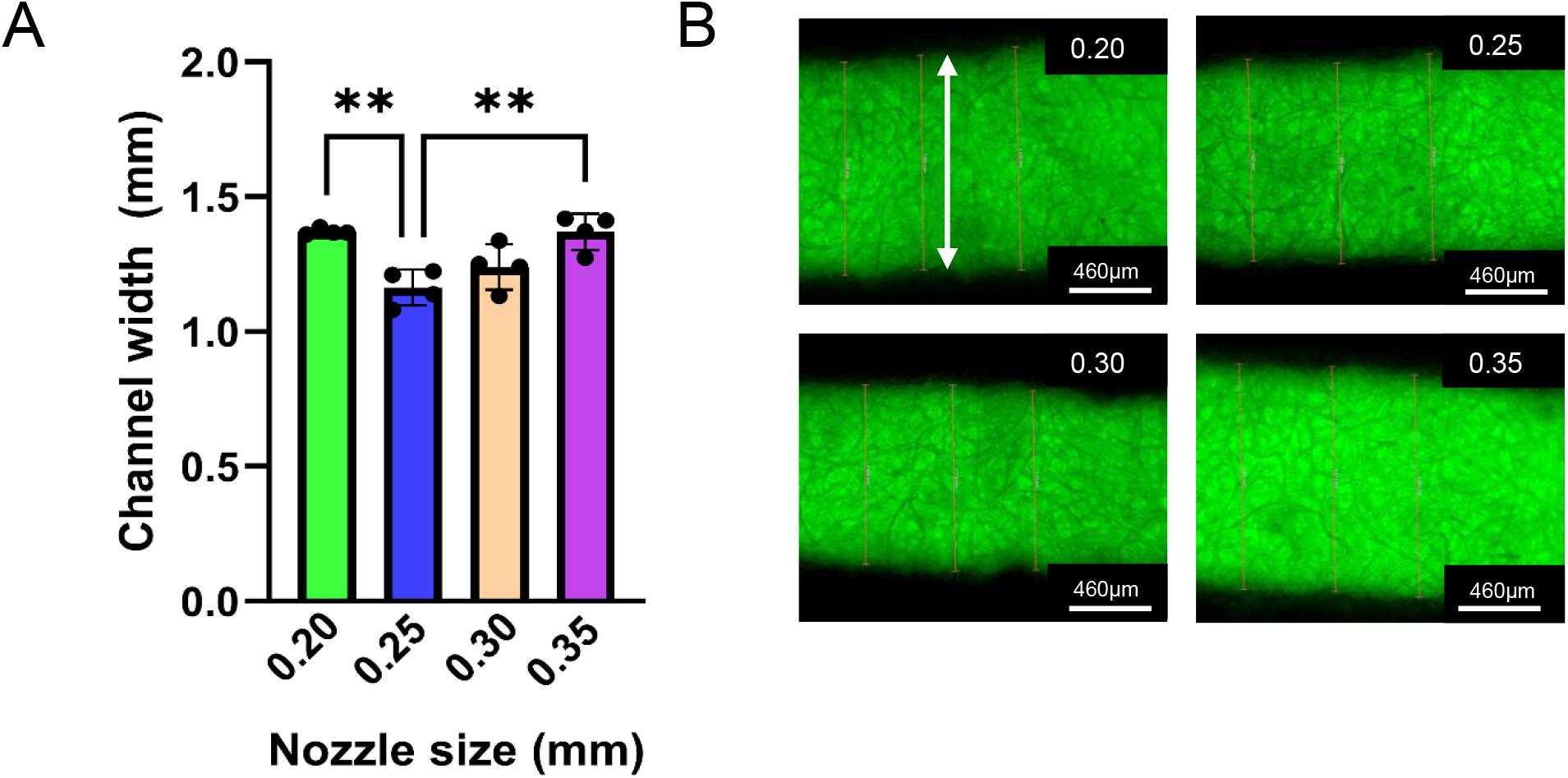
Effect of nozzle size on channel width resolution. **A**. Comparison of 1800µm PP 3D µPAD actual channel widths after thermal curing for 0.2, 0.25, 0.3 and 0.35mm nozzles (n=4 discrete devices) **B**. Fluorescence image samples for each nozzle size after deposition of ThT within the channels. **p<=0.01.

### Validation of PP 3D-µPAD with dimeric g-quadruplex formation assay

It is well established that the monomeric G-quadruplex structure binds with ThT^16^, significantly enhancing fluorescence hence rendering the complex an attractive fluorescent sensing probe for bioanalysis. In this work a G4-dimer was selected that has been proven to outperform its monomeric version. The stability of the output signal generated by the G-quadruplex structure is critical to ensuring repeatable and reliable sensing. Quadruplex formation requires the presence of monovalent cations in particular K^+^ and Na^+^ ions which exist in physiological systems^20^. The G4-dimer folds into an antiparallel topology in the presence of ThT and high concentration of Na^+^ ions^21^. The fundamental unit of the g-quadruplex is the G-tetrad composed of a planar array of 4 guanine bases associated through a network of hydrogen bonds. Na^+^ ions are located on the plane with the G-tetrad and promote the stability of the structure.

Before implementation on the PP 3D-µPAD, a solution assay was used to validate the formation of the G4-Dimer with results shown in ***Figure 5***. G4-Dimer binding with ThT in the presence of 150mM of Na^+^ enhanced the fluorescence intensity with statistical significance when compared to ThT (p < 0.001) and the control oligo by more than 100-fold (***Figure 5A***). Similarly, a comparison with varying concentrations of the control oligo and G4-Dimer, the G4-Dimer could be clearly differentiated from the control oligo with statistical significance for concentrations of 2 - 10µm (p < 0.0001) (***Figure 5B***). A lateral flow PP 3D-µPAD was designed to implement the G4-Dimer formation assay (***Figure S1***). ThT was deposited on reaction pads (2a) and (2b), stored in the dark and allowed to dry for 24 hours ***A***. To run the test, the single stranded G4-Dimer sequence was deposited on the sample pad (1a), and the control oligo was deposited on the sample pad(1b) and allowed to wick towards the test region (***Figure 6A***). On contact with (2a), ThT binding with the G4-Dimer sequence formed an anti-parallel dimeric g-quadruplex structure, which enhanced the fluorescence intensity significantly when compared to ThT binding with the control oligo. Fluorescence images captured from regions (2a) and (2b) were analyzed using Image J Fiji, the parameters of interest were the mean gray value which gave information on the average pixel intensity within the selected region of interest, skewness which provided the image’s pixel intensity distribution and the region of interest (ROI) which was the specific area that was measured. The ROI was fixed for all images to ensure consistency with the measurements. The mean gray value which directly correlates with the fluorescence intensity was plotted against the concentration of each sample and 2 way ANOVA and Sidak’s multiple comparisons test was used to compare the means between the G4-Dimer and control oligo fluorescence intensities. The G4-Dimer (2a) consistently fluoresced with a higher intensity when compared with equimolar concentrations (20-100nM) of the control oligo (2b) (***Figure 6B&C***).

**Figure 5:**
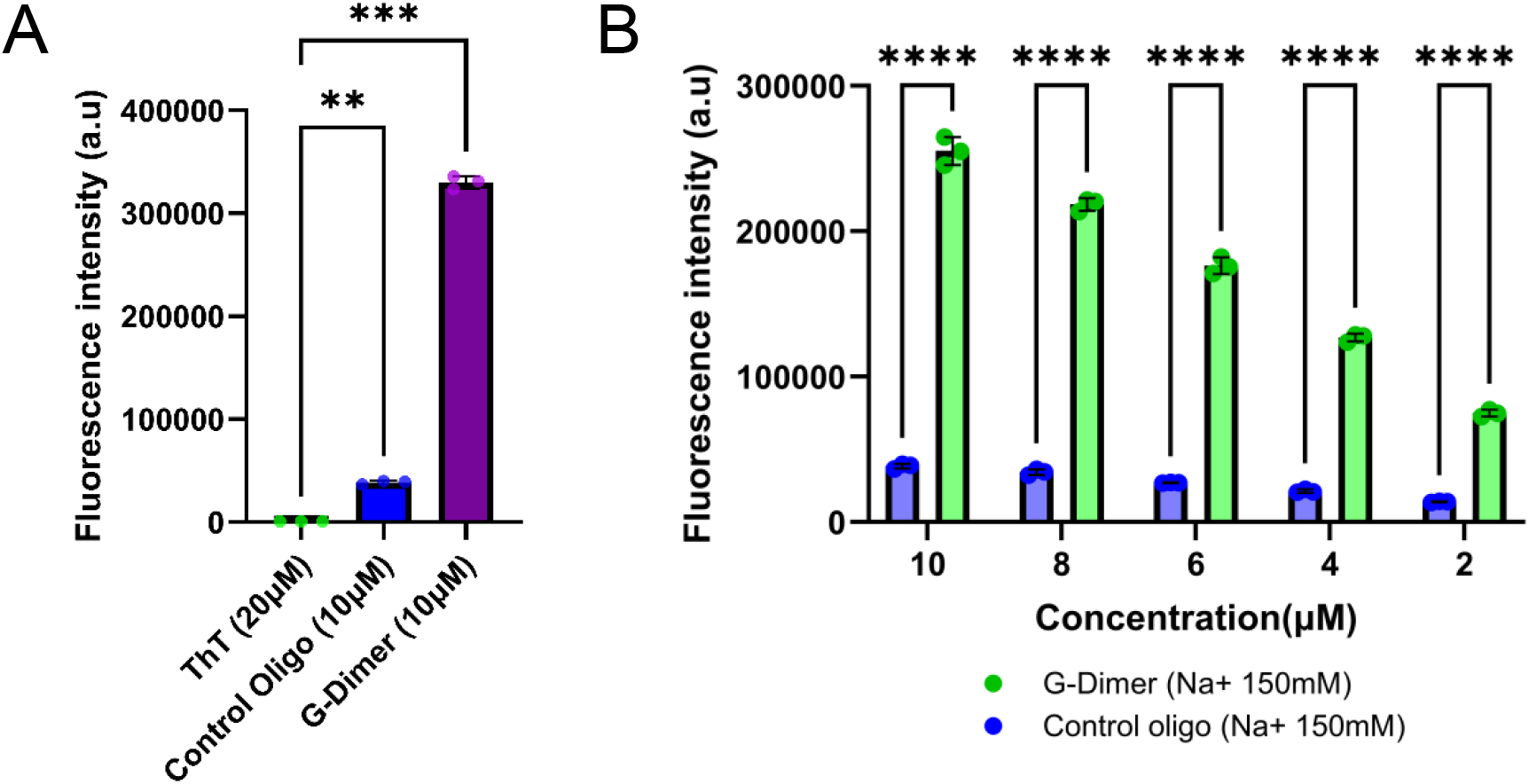
Validation of the G4-Dimer formation assay in solution. **A**. G-4 Dimer compared to baseline ThT and the control oligonucleotide **B**. Five concentrations of G4-Dimer and control oligo. **p<=0.01, ***p<=0.001, ****p<=0.0001.

**Figure 6:**
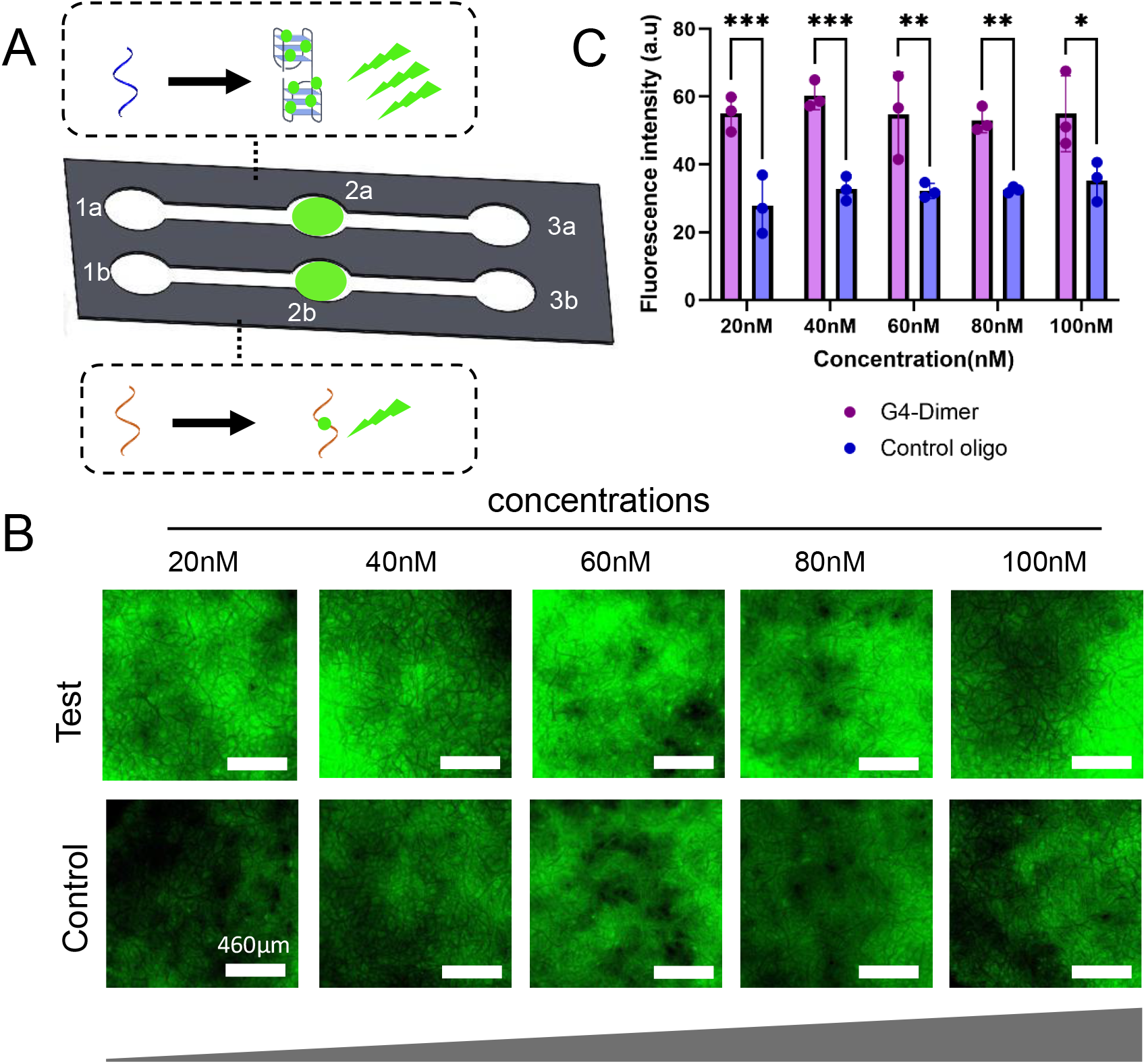
Validation of the G4-Dimer formation on paper. **A**. Schematic showing implementation of the G4-dimer formation assay, the G4 dimer ssDNA is deposited on (1a), control oligo on (1b), each will wick to the sensing region with ThT (2a and 2b) and (3a and 3b are the waste collection pads **B**. fluorescence images of each sensing pad used to quantify fluorescence intensity for five concentrations from 20nM to 100nM C. Results of fluorescence intensity quantified using Image J. *p<=0.05, **p<=0.01, ***p<=0.001

For lower concentrations (20 nM and 40 nM) p-values between the G4-Dimer and control oligo below 0.001 were obtained, however, for 60 nM and 80 nM p-values were higher (p < 0.01) with 100 nM. From the results better sensitivity was achieved with lower concentrations which can be attributed to higher signal to noise ratio and optimal stoichiometry within the paper matrix. Furthermore, at higher concentrations, excess ThT molecules can aggregate on the paper surface leading to self-quenching where excited-state energy transfers non-radiatively between stacked dyes leading lower fluorescence intensity^22^. Furthermore, these results can be attributed to the paper’s porosity composition and the degree with which the sample embeds within the paper, light scattering or uneven distribution of the sample as it wicks through the microchannels^23,24^. Despite these potential challenges, the fluorescence intensity achieved by each sample can be distinctly differentiated which demonstrates the utility of the PP 3D µPAD as a platform for the recognition of secondary nucleic acid structures.

## Conclusion

This study demonstrates the potential for 3D printing PP directly onto paper representing a transformative platform that seamlessly merges rapid prototyping precision of extrusion based additive manufacturing with paper’s inherent affordability, portability and biocompatibility. A comprehensive characterization and comparison with machinable wax, TPU and PLA demonstrated the merits of this approach and its potential to be adopted as a robust, reconfigurable and repeatable fabrication technique for the creation of high resolution paper-polymer analytical devices. Furthermore validation of the PP 3D µPAD platform using a fluorescence based assay for the formation of the G4-Dimer nucleic sensing scaffold, demonstrated its applicability for bioanalytical sensing. Future work will explore integration of the 3D printed PP open microchannels with sample processing microfluidic devices for instance serum extraction from blood, 3D printed structures to manipulate fluid flow such as valves or pumps and electrochemical sensing.

## Acknowledgements

This work was supported by a Natural Sciences and Engineering Research Council of Canada (NSERC) Vanier Canada Graduate Scholarship awarded to P.N., NSERC Discovery Grant (RGPIN-2020-04559) of K.K., Canada Foundation for Innovation John R. Evans Leaders Opportunity Fund of K.K. and R.P., University of Calgary Research Excellence Chair Award and NSERC-AI Advance grant and A-MEDICO grant of R.P.

## Data Availability

The data that support the findings of this study have been included in supplementary information and are available upon reasonable request.

## Conflict of interest

The authors declare no conflict of interest.

## Notes

### Competing Interest Statement

The authors have declared no competing interest.

