## Supplementary Information for "Extending the limits of 3D printed polymers on paper towards bioanalytical sensing"

***Table S1:*** *Raw data for 3D µPAD channel widths using four difference nozzle sizes.*

| Nozzle size | E1 | E1 Average | E2 | E2 Average | E3 | E3 Average | E4 | E4 Average |
|---|---|---|---|---|---|---|---|---|
| 3.5 | 1.407 | | 1.381 | | 1.352 | | 1.262 | |
| | 1.413 | | 1.404 | | 1.402 | | 1.306 | |
| | 1.441 | 1.420 | 1.454 | 1.413 | 1.366 | 1.373 | 1.255 | 1.274 |
| 3.0 | 1.179 | | 1.117 | | 1.323 | | 1.257 | |
| | 1.270 | | 1.127 | | 1.348 | | 1.206 | |
| | 1.270 | 1.240 | 1.151 | 1.132 | 1.339 | 1.337 | 1.291 | 1.251 |
| 2.5 | 1.200 | | 1.038 | | 1.212 | | 1.161 | |
| | 1.197 | | 1.096 | | 1.246 | | 1.115 | |
| | 1.236 | 1.211 | 1.109 | 1.081 | 1.214 | 1.224 | 1.143 | 1.140 |
| 2.0 | 1.416 | | 1.416 | | 1.35 | | 1.364 | |
| | 1.363 | | 1.332 | | 1.374 | | 1.362 | |
| | 1.354 | 1.378 | 1.354 | 1.367 | 1.378 | 1.367 | 1.356 | 1.361 |

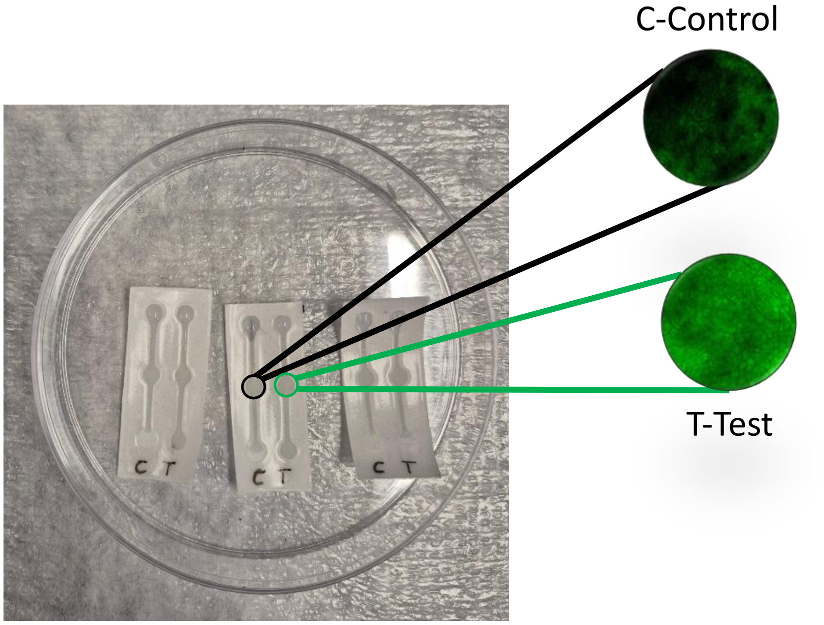


***Figure S1: Validation of g-quadruplex formation on 3D printed paper-polymer microfluidic devices.*** *n=3 devices for each concentration, C- control, T- test, control reaction pad contained a synthetic DNA sequence which exhibited lower fluorescence when compared to the test which had the g-quadruplex sequence.*

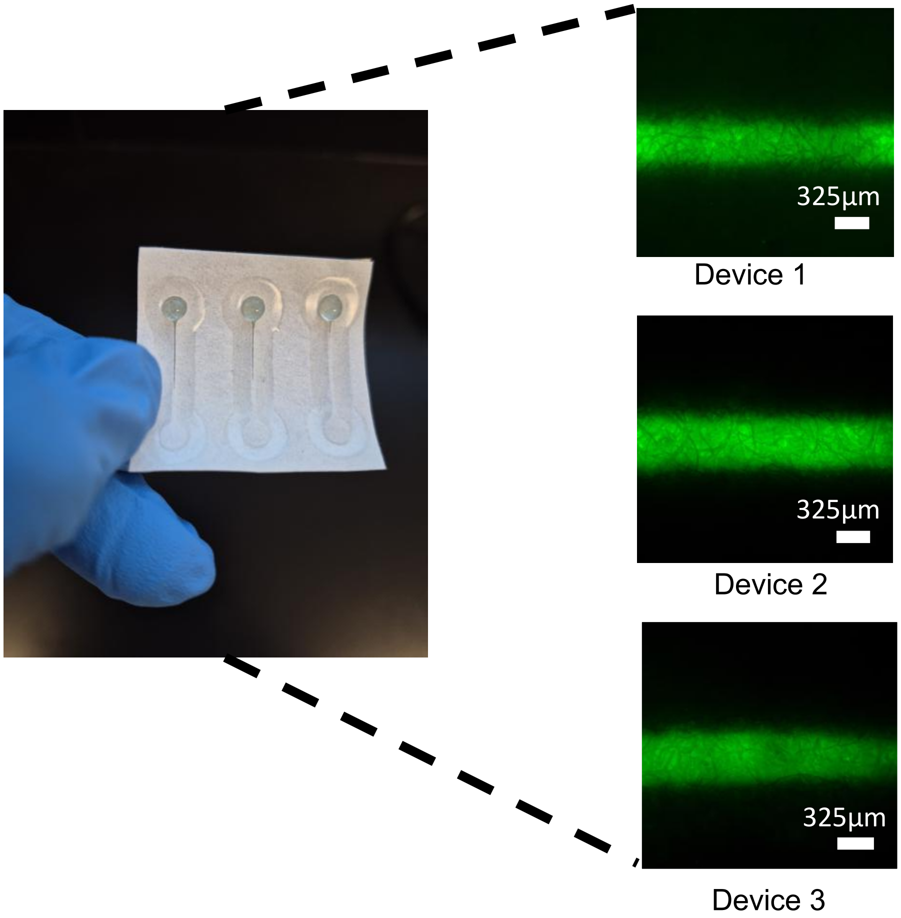


***Figure S2: Highest resolution functional channels achieved with 3D printing polypropylene directly on chromatography paper*** *(621±33μm) n=3 discrete devices were measured, results showed repeatability and robust high resolution hydrophobic barriers*

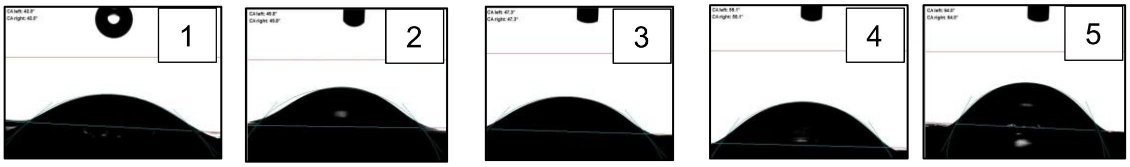


***Figure S3:*** *Contact angle measurements of 1800μm 3D μPAD device within hydrophilic microchannels n=5 discrete devices were measured (51.4 ± 8.36 ˚)*

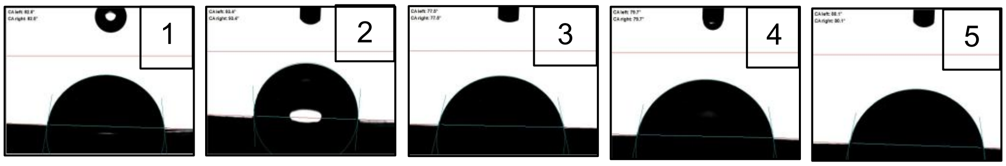


***Figure S4****: Contact angle measurements of 1800µm 3D µPAD device barriers fabricated using polypropylene n=5 discrete devices were measured ($82.6 \pm 6.27^{\circ}$)*
